# Wetland loss in the Ñeembucú Wetlands Complex, Paraguay, using remote sensing

**DOI:** 10.1101/2022.01.03.474818

**Authors:** Frances O’Leary

## Abstract

South American wetlands are of global importance, yet limited delineation and monitoring restricts informed decision-making around the drivers of wetland loss. A growing human population and increasing demand for agricultural products has driven wetland loss and degradation in the Neotropics. Understanding of wetland dynamics and land use change can be gained through wetland monitoring. The Ñeembucú Wetlands Complex is the largest wetland in Paraguay, lying within the Paraguay-Paraná-La Plata River system. This study aims to use remotely sensed data to map land cover between 2006 and 2021, quantify wetland change over the 15-year study period and thus identify land cover types vulnerable to change in the Ñeembucú Wetlands Complex. Forest, dryland vegetation, vegetated wetland and open water were identified using Random Forest supervised classifications trained on visual inspection data and field data. Annual change of −0.34, 4.95, −1.65, 0.40 was observed for forest, dryland, vegetated wetland and open water, respectively. Wetland and forest conversion is attributed to agricultural and urban expansion. With ongoing pressures on wetlands, monitoring will be a key tool for addressing change and advising decision-making around development and conservation of valuable ecosystem goods and services in the Ñeembucú Wetlands Complex.

## 1. Introduction

South American wetlands are of global importance, yet limited delineation and monitoring restricts informed decision-making around the drivers of wetland loss in the neotropics. Wetlands are some of the most valuable ecosystems on Earth, providing goods and services including water storage and purification, carbon fixation, agricultural production and provision of biodiverse habitat (Kashaigili et al., 2006; Ramsar Convention, 2016; Guo et al., 2017). Wetlands are estimated to cover around 3-6% of the Earth’s surface and South America holds a large proportion of these, with wetland area covering 20% of the continent’s surface and holding 42% of the Earth’s peat volume (Junk, 2013; Gumbricht et al., 2017; Kandus et al., 2018).

Despite their value, wetlands are one of the Earth’s most vulnerable ecosystems (Millennium Ecosystem Assessment, 2005). Wetlands are being lost at a faster rate than any other ecosystem, with over half of Earth’s wetlands becoming degraded or lost in the last 150 years (Sica et al., 2016; Slagter et al., 2020). Wetland degradation, destruction and modification has been driven by anthropogenic and natural pressures (Baker et al., 2007; Gardner et al., 2015; Reis et al., 2017). In South America, a growing human population and increasing demand for agricultural products has driven infrastructure development, agricultural expansion and exploitation of natural resources, exerting pressure on wetlands. Extreme weather events such as drought and storms can also drive wetland change. Wetland destruction and degradation reduces the capacity of wetlands to provide valuable ecosystem services, including reduced flood and drought mitigation, wetland biodiversity loss, and reduced provisioning of natural resources.

Despite global concern for wetland habitats and the ecosystem goods and services they provide, little is known about the extent of wetland conversion in South America (Junk, 2013). Paraguay is one of South America’s least studied countries, and even less is known about the largest wetland within Paraguay’s administrative boundaries, the Ñeembucú Wetlands Complex (Kandus et al., 2018; Pett and Wyer, 2020; Rosset et al., 2020). The Ñeembucú Wetlands Complex lies within the Paraguay-Paraná-La Plata River system, which has the 9^th^ highest water discharge into oceans and 5^th^ highest drainage area of rivers worldwide (Milliman and Meade, 1983; Junk, 2013). The Paraguay-Paraná-La Plata River system flows from tropical to temperate regions, resulting in high environmental heterogeneity and biodiversity (Sica et al., 2016). The Ñeembucú Wetlands Complex has a humid subtropical climate, with 1604mm average total annual precipitation and follows a dry/wet season trend (Beck et al., 2018; Climate-Data, 2021). Ñeembucú is the 3^rd^ least populated department in Paraguay, and livelihoods within this department predominantly rely on agriculture and local fisheries (UNFPA and DGEEC, 2021). Local populations are dependent on the ecosystem health of the wetlands as a result. Common agricultural practices in the area include the use of fire to promote growth of palatable grasses and subsistence deforestation.

Utilisation of monitoring to understand wetland dynamics and land use change trends is crucial for effectively informing decision-making and development planning in the Ñeembucú Wetlands Complex. Wetland monitoring is important for assessing global change, identifying areas at high risk of land conversion and degradation, and examining the effectiveness of policy in preserving wetland habitats (Lang and McCarty, 2008; Dewan and Yamaguchi, 2009). Knowledge of the pace and extent of wetland change can be gained using remote sensing techniques and this understanding is required to effectively manage wetland resources and development. Remote sensing enables studies on greater spatial and temporal scales and is less expensive than field studies (Kandus et al., 2018). However, wetland monitoring using remote sensing has faced challenges as wetland habitats are highly variable and lack unifying features which enable identification (Gallant, 2015). Recent developments in remote sensing technology have allowed advancements in wetland delineation, mapping and monitoring with high-quality, high-resolution satellite imagery (Junk, 2013). In particular, optical data is limited in its ability to detect hydrology and a shift in data sources for wetland mapping to synthetic aperture radar data has been seen with the availability of the Sentinel-1 collection (Guo et al., 2017). Recent developments can be utilised to gain knowledge about wetland dynamics in the Ñeembucú Wetlands Complex.

The objectives of this study are to use remotely sensed data to map land cover between 2006 and 2021, use these maps to quantify wetland change over the 15-year study period and thus identify land cover types vulnerable to change in the Ñeembucú Wetlands Complex.

## 2. Materials and Methods

Land cover was identified and quantified for a series of years within a 15-year period from 2006 to 2021. A two-step classification process was followed; step 1 identified forest, non-forest vegetation, and open water cover, and step 2 identified dryland vegetation and wetland vegetation, differentiated within the vegetation class from step 1.

Forest was defined using the national forest definition, characterised by the presence of trees and at least 10% canopy cover (FAO, 2020). Vegetated wetland included all seasonal and permanent wetland types except open water, and this primarily consisted of freshwater marshes, peatland, seasonally-inundated grassland and shrub-dominated wetland in the study area (Ramsar, 1990). Dryland was dry vegetation with less than 10% canopy cover from trees, and open water was areas of water not covered by vegetation.

Supervised classifications were carried out for four ‘supervision years’; 2006, 2011, 2016 and 2021. Three further ‘intermediate years’ (2009, 2014 and 2019) were classified using the classifier trained on the nearest ‘supervision year’. The area covered by each land cover class was quantified for each year and change over the study period was measured.

### 2.1 Study Area

The study area is an 8,361km^2^ region within the Department of Ñeembucú, Paraguay (see Figure 1). The study area is bordered by the River Tebicuary in the north, River Paraguay in the west, River Parana in the south and the department’s administrative boundary in the east. The Ñeembucú Wetlands Complex is located at the confluence of two of South America’s most important rivers, the Paraguay and the Paraná, and is part of the Rio de la Plato Basin System (ymin: −27.44417, ymax: −26.39394, xmin: - 58.66491, xmax: −57.18209). Ñeembucú has the 3^rd^ lowest population size of Paraguay’s departments (UNFPA and DGEEC, 2021).

**Figure 1.**
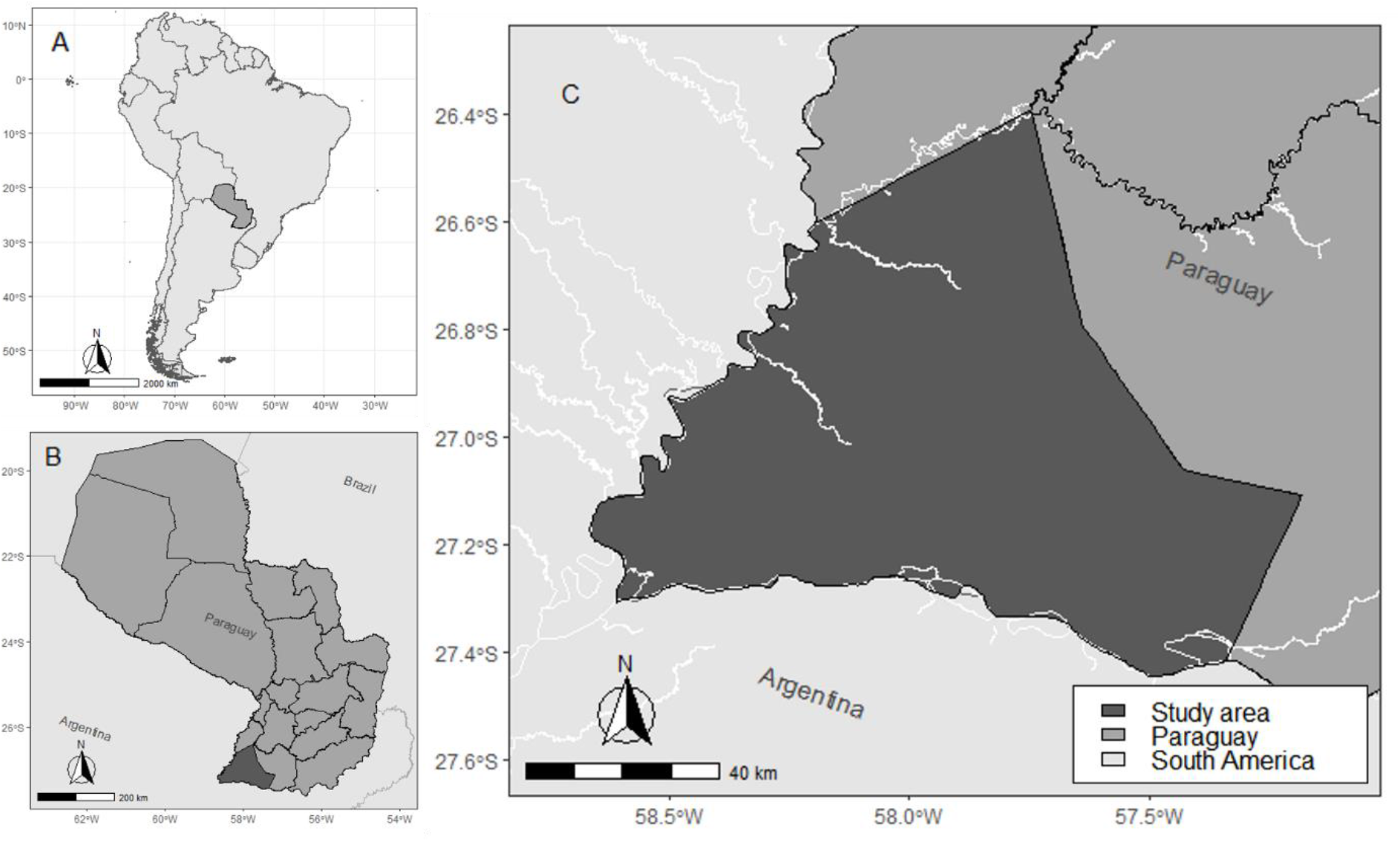
A study area map showing; **A** the location of Paraguay within South America, **B** the location of the study area within Paraguay, and **C** the study area locally. Maps were created using RStudio, and the Paraguay administrative boundaries were sourced from UNFPA and DGEEC (2021) (R Core Team, 2021).

### 2.2 Datasets

#### 2.2.1 Validation Data

##### Field Data

Field data was collected in October 2021 from 129 plots within 11 localities across the study area (See Figure 2). The localities were selected due to being either being public access land, properties for sale with surveying permissions from the owner, or properties that we had previously established relationships and permission to survey the property. Random allocation of plots for field data collection was not feasible for the study area due to the high proportion of privately owned land (Fian International, 2021). Between 5 and 21 plots were visited at each locality, depending on locality size. Each plot is 10m^2^ and the plots were distributed evenly across habitat types in each locality and at least 100m apart. Each plot was recorded as dryland, vegetated wetland or forest. The forest plots were excluded from the dataset and 97 sample plots remained as field data to be used in producing a non-forest vegetation classification. The dataset was randomly split into a training partition (75% of observations) and a testing partition (25% of observations).

**Figure 2.**
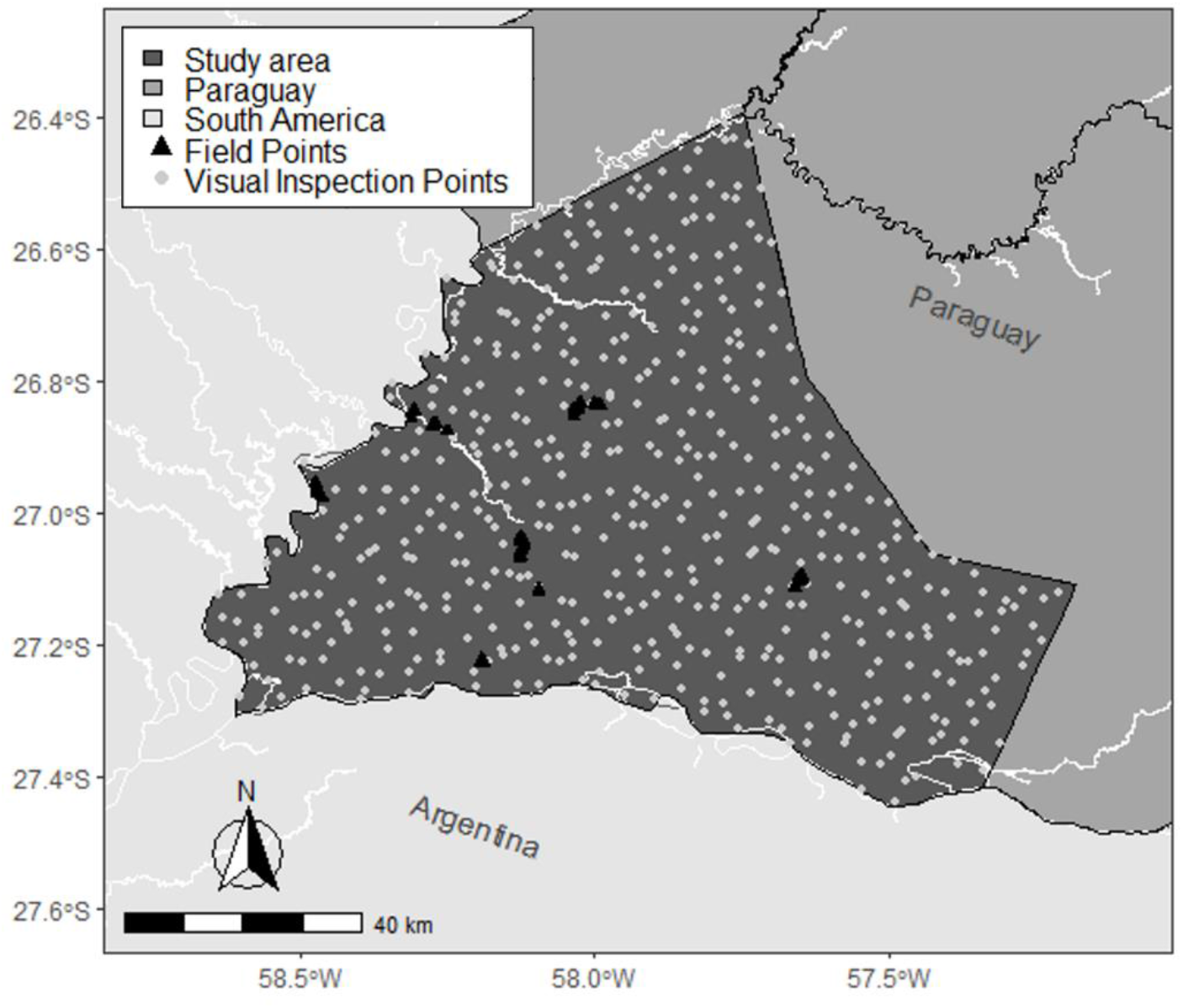
A map showing the distribution of ground-truth data used to supervise the land cover classifications. Both groups of points were split randomly into a 75% training partition and 25% testing partition. The visual inspection points (n = 504) supervised the Level 1 classification and the field points (n = 129) supervised the Level 2 classification.

##### Visual Inspection Data

Image interpretation data was collected using Sentinel and Landsat images from October in supervision years: 2006, 2011, 2016 and 2021 (Copernicus, 2021; USGS, 2021). This was done by visually inspecting 504, randomly allocated, 30m^2^ plots on true colour image composites from the available satellite imagery for October of that year. Resolutions ranged between 10m^2^ and 30m^2^ (See Figure 2). The visual inspection plots were allocated across the study area using random stratified sampling using the ‘sp’ package v.1.4 in RStudio version 3.7.2 (Pebesma and Bivand, 2005; R Core Team, 2021). For 2006 and 2011, Landsat 7 Surface Reflectance and Landsat 5 Surface Reflectance were used to create composite images for inspection. For 2016 and 2021, Sentinel-2 Surface Reflectance and Landsat 8 Surface Reflectance were used to create composite images for inspection. One composite image from each dataset was produced for each year, and every sample point was inspected and identified as forest, non-forest vegetation or open water based on the plot’s appearance. Forest appeared dark green, open water appeared black or blue, and non-forest vegetation belonged to neither of the aforementioned classes. Vegetated wetland and dryland could not be differentiated through a visual inspection because of the high variability and inconsistency in appearance (Kandus et al., 2018). The visual inspection dataset was randomly split into a training partition (75% of observations) and a testing partition (25% of observations).

#### 2.2.2 Classification Data

Images from the USGS Landsat Collections (Landsat 5, 7 and 8, Level 2 [Collection 2]) and Sentinel Collections (Sentinel-1 SAR GRD and Sentinel-2 MSI) were sourced using Google Earth Engine for the study period between 2006 and 2021 (Gorelick et al., 2017; Copernicus, 2021; USGS, 2021).

### 2.3 Identification of land cover

Imagery from the LANDSAT and Sentinel missions were utilised to develop supervised classifications of land cover over the study period (Copernicus, 2021; USGS, 2021). A two-step methodology was employed to firstly identify open water, vegetation, and forest (Level 1 classification), and secondly to identify dryland and vegetated wetland within the vegetation class (Level 2 classification).

#### 2.3.1 Data processing

Imagery from the Landsat and Sentinel missions over a 12-week period (23^rd^ August – 14^th^ November) were used to calculate Enhanced Vegetation Index (EVI), Normalized Difference Vegetation Index (NDVI), Normalized Difference Water Index (NDWI) from Landsat data and the 10^th^ percentile, 90^th^ percentile and difference between the 10^th^ and 90^th^ percentile for Sentinel-1 bands. The annual seasonality of NDVI, NDWI and Bare Soil Index (BSI) were calculated over the year leading up to the end date of the 12-week imagery period. The mean value of bands in each pixel were used to produce a composite image of the study area. SRTM Digital Elevation Data was also collated at 90m^2^ and a mean taken for each 30m^2^ pixel of the composite image (Jarvis et al., 2008). The final composites for each year contained raw bands and processed bands (see Table 1). The equations used in band processing are as follows;

**Table 1.**
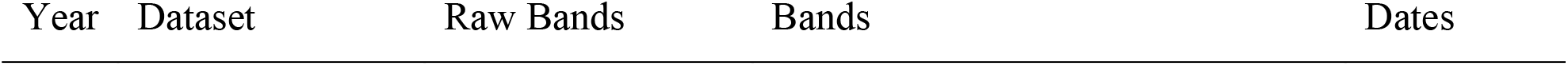
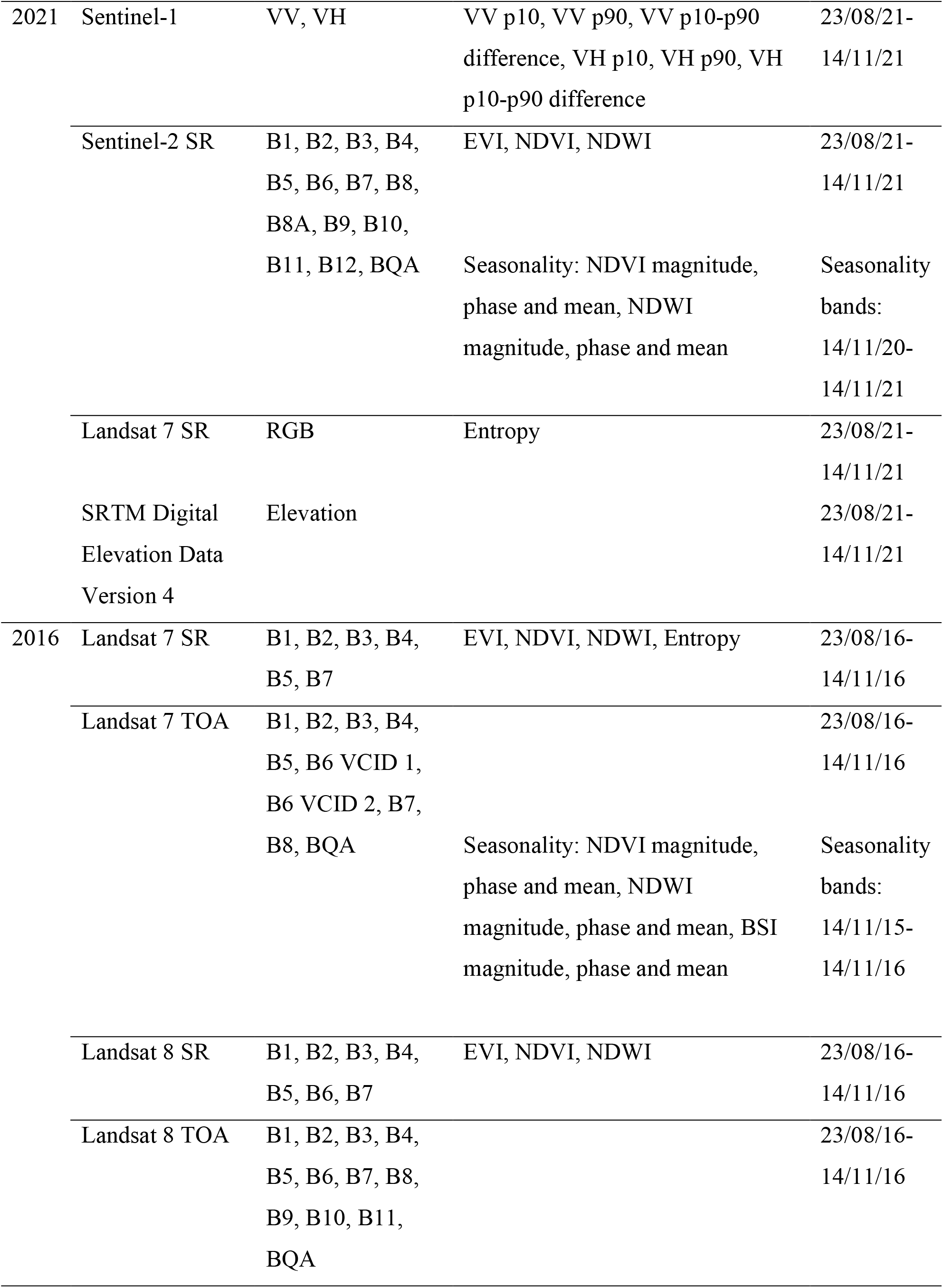
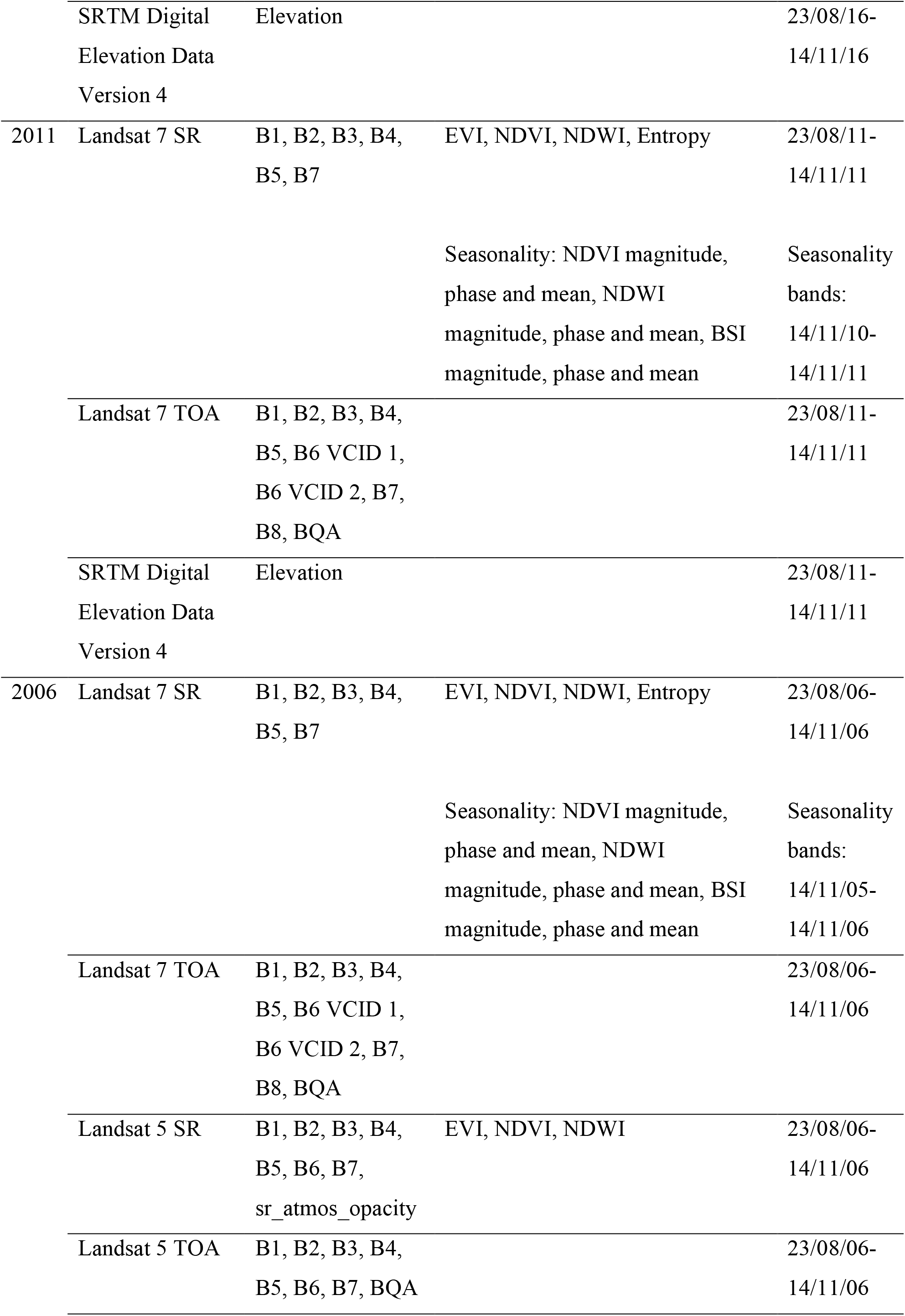
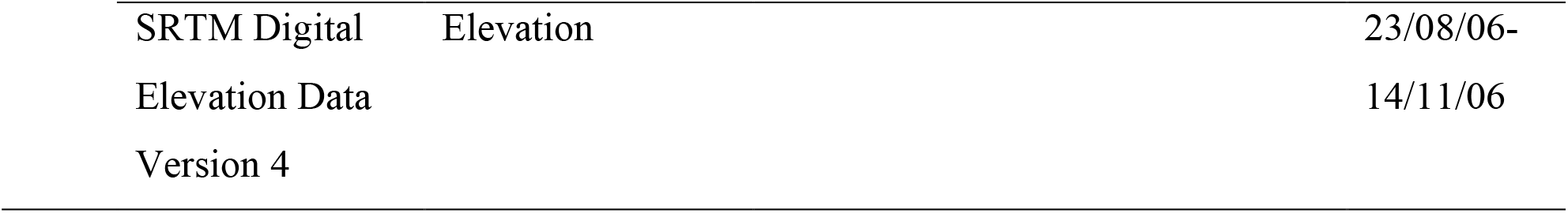
Datasets and image bands used in the classification of land cover.

For NDVI:

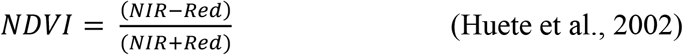

For NDWI:

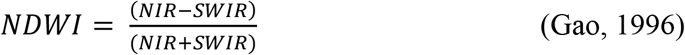

For BSI:

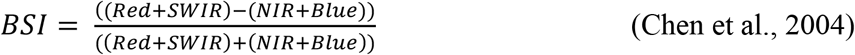

For EVI:

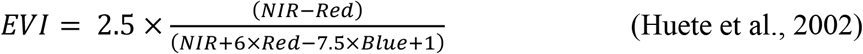

#### 2.3.2 Classification

For the Level 1 classification, which identified open water, vegetation, and forest, a random forest classifier, with 800 trees, a minimum leaf population of 1 and a bag fraction of 0.5, was trained using the training partition of visual inspection data. A supervised classification was performed for each study year (2006, 2011, 2016, 2021), creating a classification of forest, vegetation and open water for each year. In order to produce a Level 1 classification for three intermediate years (2009, 2014, 2019), the trained classifier of the closest year was used to classify the composite image of the intermediate year. For example, a classification for 2009 was produced using the classifier trained on 2011 data. A classification for 2014 used the 2016-trained classifier and a classification for 2019 used the 2021-trained classifier. This produced a classification of forest, vegetation, and open water for 7 study years between 2006 and 2021. There was no satellite imagery available for 35.8km^2^, 0.4% of the study area, in the 2021 study period. These pixels were assumed not to have changed since 2020 and were assigned values from the 2020 classification.

For the Level 2 classification, identifying dryland and vegetated wetland, a random forest classification, with 400 trees, a minimum leaf population of 1 and a bag fraction of 0.75, was trained using the training partition of the field data. A supervised classification was performed on the area classified as vegetation in the Level 1 classification for each study year (2006, 2011, 2016, 2021), creating a vegetation type classification. The same three intermediate years classified in the Level 1 classification underwent vegetation type classification.

The Level 2 classification was combined with the Level 1 classification image for each study year, producing classified land cover images showing forest, non-forest dryland, vegetated wetland and open water. Land cover classifications are in 30m^2^ resolution and use a WGS84 coordinate reference system.

#### 2.3.3 Accuracy assessment

To assess the accuracy of each classification, a confusion matrix carried out on Google Earth Engine (Stehman, 1997; Gorelick et al., 2017). A further accuracy assessment was carried out in RStudio, which quantified the total accuracy of each classification and the number of false positives and number of false negatives for each class (R Core Team, 2021).

Accuracy assessments were carried out for the land cover maps in supervision years, which were 2006, 2011, 2016, and 2021. These were carried out on the Level 1 classification outputs and Level 2 classification outputs separately.

### 2.4 Change Detection

The area covered by each land cover class was measured from the land cover classification for each study year and the mean annual change between each of the images was calculated. The mean annual change for the whole study period was calculated by taking the mean of annual change estimates between study years.

### 2.5 Precipitation Trend

Precipitation data was sourced from the CHIRPS daily (version 2.0) climate dataset at 5566m resolution (Funk et al., 2015). The total annual precipitation was quantified by taking the sum of total annual precipitation of all pixels across the study area for every year between 2006 and 2021 on Google Earth Engine (Gorelick et al., 2017). The total annual precipitation was then plotted using RStudio (R Core Team, 2021).

## 3. Results

### 3.1 Land Cover Classification

Classified land cover maps of the study area, produced in the two-step classification methodology, are presented in Figure 3. The Ñeembucú Wetlands Complex is dominated by vegetated wetland, covering 65-79% of the study area between 2006 and 2021. Second to vegetated wetland was dryland, covering 8-23% of the study area over the study period. In 2016, vegetated wetland covered 11% more of the study area than for the same class in 2014. Inversely, dryland covered 11% less of the study area than for the same class in 2016. This is explained by severe flooding due to repeated heavy rainfall in 2015-2016, which was reported in the Paraguay River basin (Dos-Gollin et al., 2018). After excluding 2016 observations due to extreme weather, vegetated wetland and dryland covered 65-70% and 19-23% of the study area, respectively, between 2006 and 2021. Forest and open water covered 6-8% and 5-7% of the study area, respectively, over the study period.

**Figure 3.**
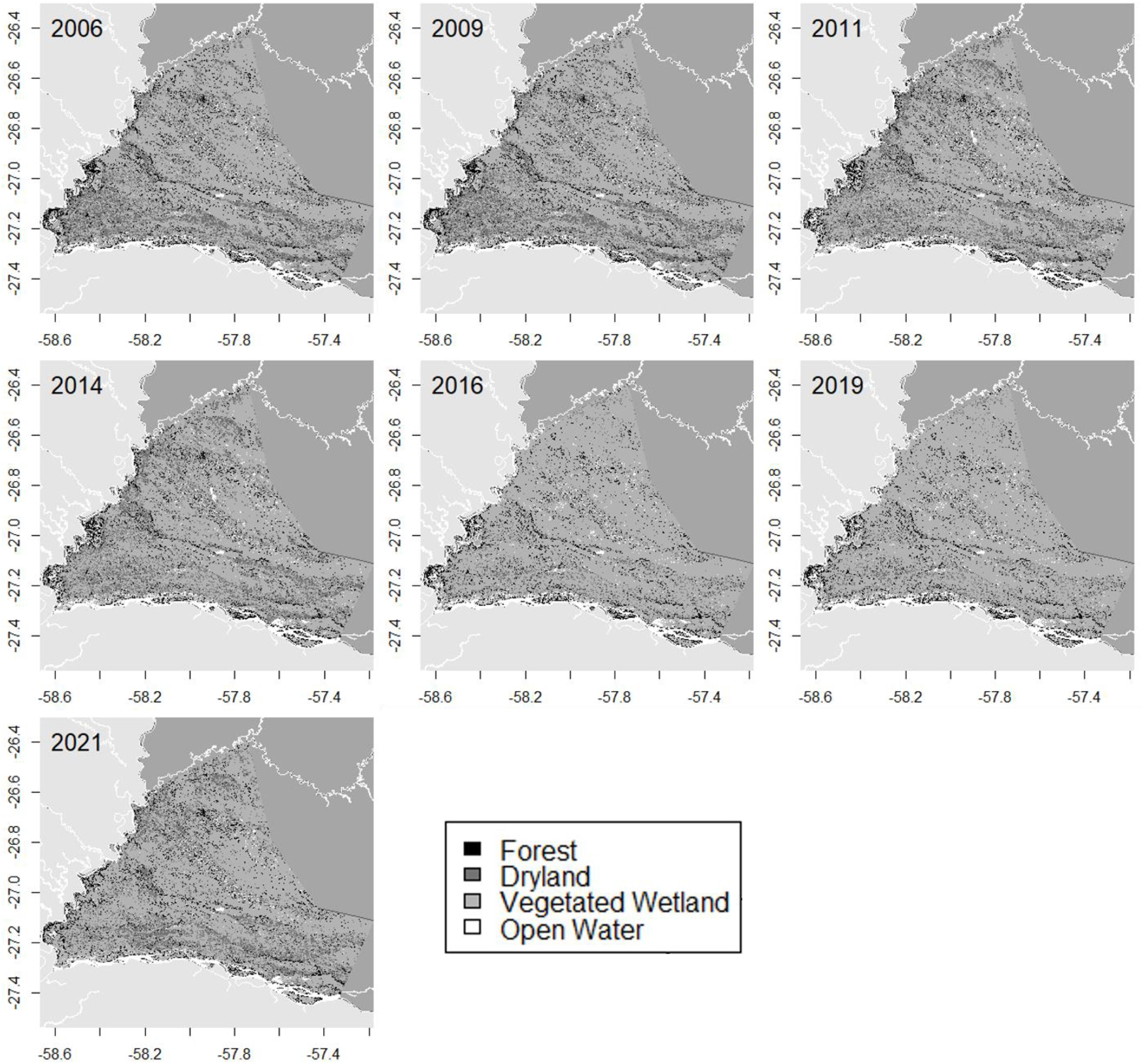
Classified land cover maps of the study area for study years between 2006 and 2021. The supervised classification years were 2006, 2011, 2016 and 2021. 2009, 2014 and 2019 were intermediate years, classified by the closest years classifier. Classifications created on Google Earth Engine and plotted in RStudio (Gorelick et al., 2017; R Core Team, 2021).

### 3.2 Classification Accuracy Assessment

The random forest classification produced Level 1 land cover classification maps with 91-96% overall accuracies and Level 2 land cover classification maps with an 82% overall accuracy. Table 2 presents the overall accuracy, and percentage of false negatives (Type I errors) and false positives (Type II errors) in each land cover class in supervision years (the years in which training and testing data were available to supervise classifications).

**Table 2.**
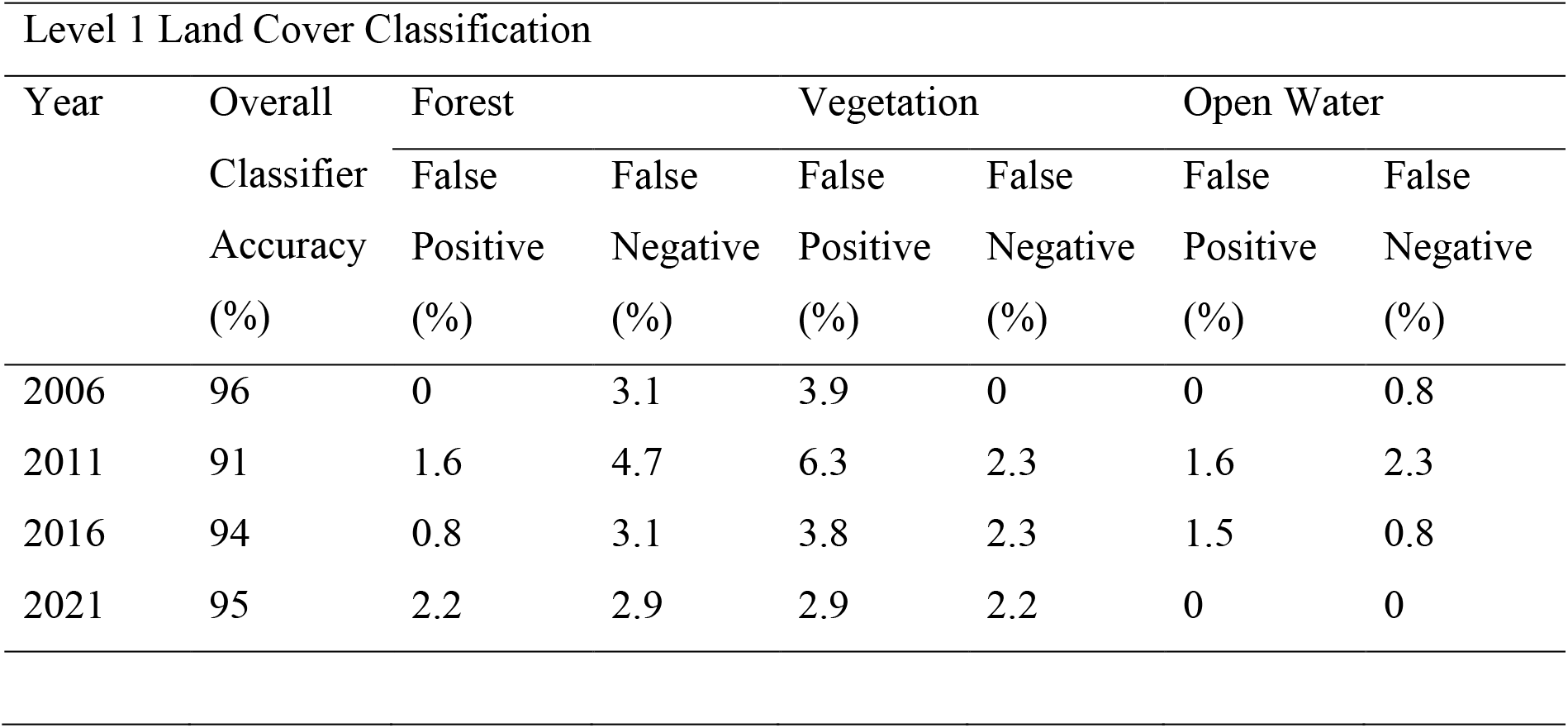
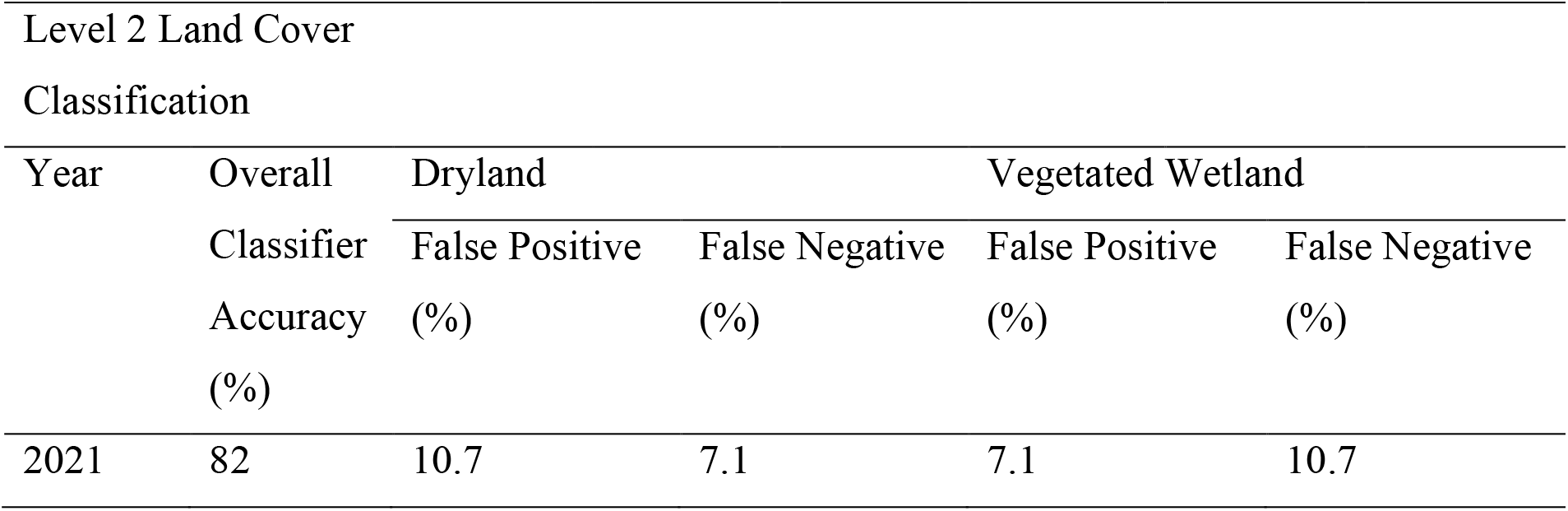
The accuracy of each classification in identifying each land cover.

Within the Level 1 Classification, vegetation was overrepresented for all years. Forest was the most underrepresented class in all years, with false negatives ranging between 2.9% of test observations in 2021 and 4.7% in 2011. In 2006, 0.8% and 3.1% of open water and forest observations, respectively, were falsely identified as vegetation. In 2011, over two thirds of the vegetation false positives were classified as forest in the testing data, and the remaining were open water. The proportion of vegetation false positive belonging to forest and open water was similar to in 2011, with over two thirds of the vegetation false positives classified as forest in the testing data. In 2021, no errors were identified in the open water class, and there was a 0.7% greater false positive identification of vegetation than of forest.

The Level 2 classification had a lower accuracy than the Level 1 classification, due to the heterogeneity in habitat types within non-forest vegetation leading to a lack of unifying features within classes (Gallant, 2015). Within both dry and wetland vegetation, dominance of grasses, herbaceous plants and shrubs vary, and the seasonality of water presence varies within the vegetated wetland class too. Within the Level 2 classification, dryland was overrepresented, with a greater number of false positives than the vegetated wetland class.

### 3.3 Land Cover Change Detection

The greatest annual change throughout the study period was observed in the dryland vegetation and vegetated wetland land cover classes (see Table 3). Extreme change in dryland and vegetated wetland was seen between 2014 and 2016, with changes of - 65.76% and 7.72% observed in each class, respectively. The extreme figures observed in 2016 are the results of extreme weather, and this year’s classification was removed from the change detection as a result (Dos-Gollin et al., 2018; Figure 4). The change detection showed vegetated wetlands decreasing at a mean annual rate of 1.65%, and a mean annual increase in dryland of 4.94% (Table 3). Further to this, forest is lost at a rate of 0.34% annually, while open water is gained at a mean rate of 0.40% annually.

**Table 3.** Mean annual change (%) in area in each land cover between study years.

| Time Period | Forest | Dryland | Vegetated Wetland | Open Water |
| --- | --- | --- | --- | --- |
| 2006-2009 | -9.12 | 3.89 | -0.92 | 3.37 |
| 2009-2011 | 9.00 | -9.69 | 1.80 | 1.34 |
| 2011-2014 | 3.54 | -1.13 | -0.52 | 4.42 |
| 2014-2019 | -3.70 | 1.54 | 0.13 | -3.07 |
| 2019-2021 | -1.43 | 30.07 | -8.74 | -4.07 |
| Overall Study Period | -0.34 | 4.94 | -1.65 | 0.40 |

**Figure 4.**
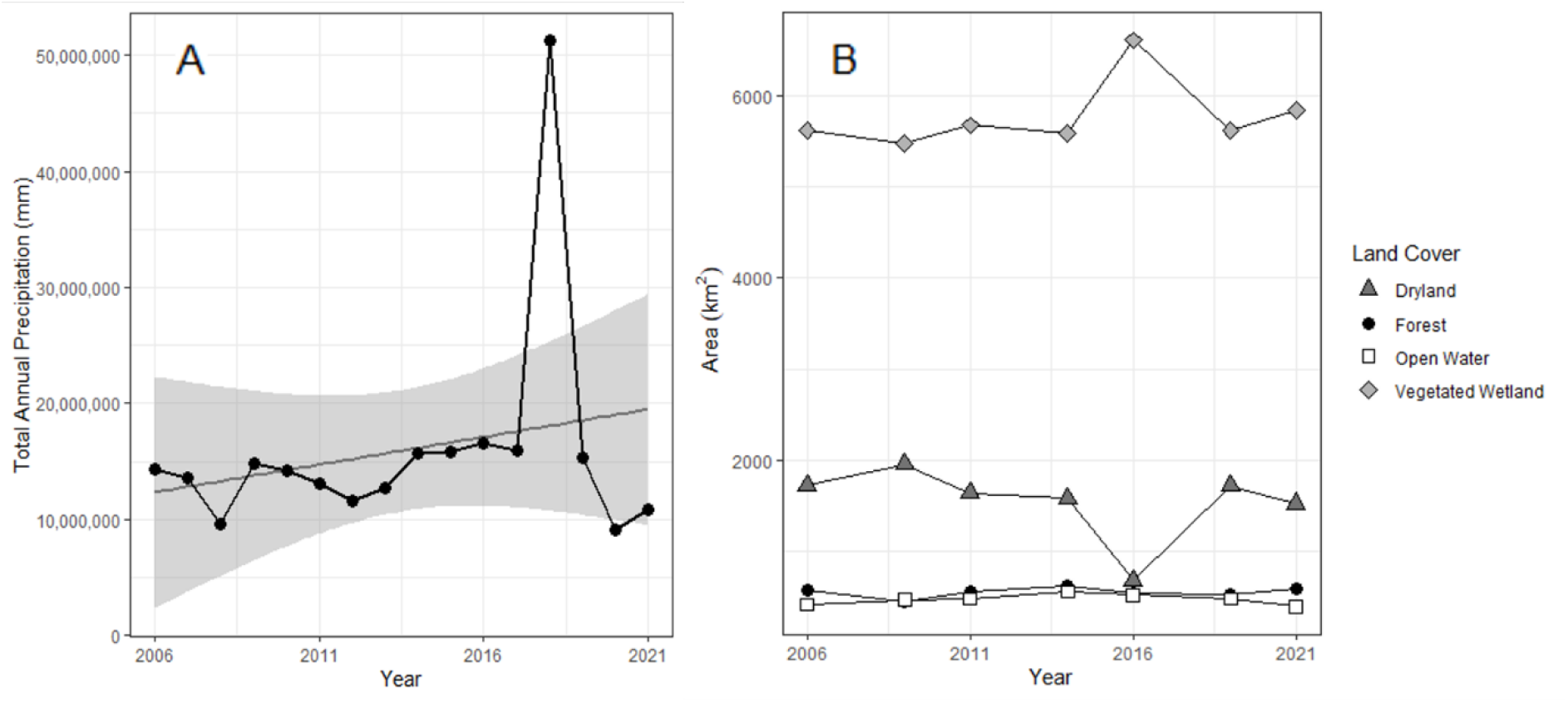
Temporal pattern of **A** total annual precipitation (mm) and **B** area (km^2^) under each land cover class in the study area between 2006 and 2021. In **A**, the observed annual precipitation is plotted in black and the fitted precipitation trend in grey. Sources: The CHIRPS daily (version 2.0) climate dataset (Funk et al., 2015). Processed in Google Earth Engine and plotted in RStudio (Gorelick et al., 2017; R Core Team, 2021).

### 3.4 Precipitation Trend

Total annual precipitation ranged between 9,036,984mm and 16,546,476mm and displayed an increasing trend over the study period (see Figure 4). However, greater variability in annual precipitation is seen in the latter years within the study period, with total annual precipitation more than three times greater than the previous year observed in 2016, and some of the lowest rainfall years observed in 2020 and 2021 (Figure 4a). The impact of extreme precipitation in 2016 is observed in the inundation of a greater area of non-forested land in that year (Figure 4b).

## 4. Discussion

The wetland change identified in the Ñeembucú Wetlands Complex is comparable to wetland change reported in regions of the Paraguay-Paraná-La Plata River system, where pressure from human activities events is driving wetland conversion and degradation trends (Collischonn et al., 2001; Junk, 2013). In the Lower Paraná River Delta, one third of freshwater marshes were converted to pasture and forestry between 1999 and 2013 (Sica et al., 2016). Similarly, Guerra et al. (2020) projected a 3% loss in native vegetation by 2050 in the Pantanal, the lowland region of the Upper Paraguay River Basin. Brandolin et al. (2013) found a 15% loss in flooded area in Córdoba, Argentina, and area in which agricultural expansion has driven high channelisation of the wetlands between 1987 and 2007. Conversely, a 66% increase in flooded area was seen in Santa Fe, a region experiencing lower agricultural pressure, within the same study in Argentina. The increase in flooded area observed in Santa Fe was attributed to increased flooding driving expansion of wetlands in a region with low agricultural pressure (Brandolin et al., 2013). The findings of this study suggest that wetland areas within the Ñeembucú Wetlands Complex are being converted to dryland, a similar trend observed in other regions within the Paraguay-Paraná-La Plata River system, and globally (Kashaigili et al., 2006; Junk, 2013; Gardner et al., 2015).

Land use in the Ñeembucú Wetlands Complex is predominantly agricultural and it is likely that agricultural and urban expansion is driving the drainage and conversion of wetlands to dryer, productive lands (Bucher & Huszar, 1995; JICA-CEPAL, 2013). This trend is seen in wetlands both globally and within the Paraguay-Paraná-La Plata River system. Wetland loss in the Ñeembucú Wetlands Complex is comparable to that seen in Argentina, where the use of water management infrastructure, such as channels and levees, has been held responsible for driving wetland conversion (Bucher and Huszar, 1995; Brandolin et al., 2013; Sica et al., 2016). In wetlands with high agricultural production in the Paraná River Delta in Argentina, artificial drainage channels were constructed to mitigate the impacts of frequent flooding caused by an increasing rainfall trend in the latter half of the 20^th^ century, and illegal construction of channels by landowners followed (Brandolin et al., 2013). Within the Lower Paraná River Delta, water management practices, cattle density, and accessibility were the primary drivers of wetland conversion (Brandolin et al., 2013; Sica et al., 2016). In the Upper Paraguay River Basin, native vegetation loss was driven by commodity agriculture, protection status, and accessibility (Guerra et al., 2020). The relative influences of these variables differed spatially, with agriculture having a lesser effect and distance to roads having a greater effect in the Pantanal wetlands compared to the dryer surrounding Cerrado and Amazon biomes.

Wetland conversion observed in the Ñeembucú Wetlands Complex is likely not attributed to precipitation, as total annual precipitation and extreme precipitation trends are increasing in the region. Doyle and Barros (2011) found increasing precipitation localised to both the Middle Paraná and Middle Paraguay Basins, in which the Ñeembucú Wetlands Complex lies, and Haylock et al. (2006) reported increased annual precipitation and extreme precipitation days, with a shortened wet season, for Paraguay and the surrounding region. Further to this, an increasing trend was seen for total annual precipitation in the Ñeembucú Wetlands Complex, within this study. Wetland dynamics are largely driven by precipitation, and without simultaneous urban and agricultural development, increasing precipitation is expected to drive greater inundation and flooding (Collischonn et al., 2001; Prieto, 2007; Pereira et al., 2021). The aforementioned increasing inundation observed in Santa Fe, Argentina, is an example of wetland expansion driven by increasing precipitation (Brandolin et al., 2013). The precipitation trends observed in the study area and the surrounding region suggest climate change is not driving conversion of wetlands observed in this study.

Continued loss of vegetated wetlands and forest in the Ñeembucú Wetlands Complex will reduce the capacity of the ecosystem to provide valuable goods and services, including water storage, provisioning of fish and fuel, and supporting wetland biodiversity. Recent developments in the region including the Coastal Defences of Pilar and Alberdi-Pilar Ruta constructions pose a further threat this vulnerable habitat (Gardner et al., 2015; MOPC, 2021a; MOPC, 2021b). The primary goals of these developments are to alleviate flood risk and increasing accessibility to Ñeembucú’s main city, Pilar, which are frequently acknowledged as drivers of wetland conversion. Further to this, development of the floodplain in Ñeembucú may reduce water storage and drive flooding in the rest of the region (Gottgens et al., 2001). It may also be the case that land use change and river modifications upstream of the Ñeembucú Wetlands Complex are influencing wetland change by moderating river discharges (da Silva and Girard, 2004). Given the clear impact global change has already had on these wetlands, wetland monitoring is an essential tool for preserving the economic, ecological and cultural value of the Ñeembucú Wetlands Complex (Sica et al., 2016; Kandus et al., 2018; Guerra et al., 2020).

Continued monitoring of the Ñeembucú Wetlands Complex and further analysis of the drivers of land use change in the region are essential for well-informed decision-making in the region (Junk, 2013; Guo et al., 2017; Kaplan and Avdan, 2018). Globally, the value of wetlands has rarely been seriously considered within decision-making (Woodward and Wui, 2001). However, integration of the value of wetlands into decision-making and development-planning will promote conservation of economically, ecologically, and culturally valuable wetland habitats. Further analysis of change within wetland types, and the drivers of this change, will be essential for identifying vulnerable habitats, monitoring wetland health and understanding the role of policy and development in driving wetland dynamics in Ñeembucú (Gumbricht et al., 2017; Davidson & Finlayson, 2018).

## 5. Conclusion

With around 6000km^2^ of wetland area within an 8000km^2^ complex of forest, grassland and wetland, the Ñeembucú Wetlands Complex is a valuable region within the Paraguay-Paraná-La Plata River system within which preservation of biodiversity, provisioning of natural resources, and water storage must be considered within development process. Within the Ñeembucú Wetlands Complex, vegetated wetlands and forest have been lost over the last 15 year, predominantly being converted into more productive, dryland areas. Given the increasing precipitation trends identified in the region, it’s likely that agricultural and urban development is driving land use change in the region. With large, ongoing, developments in the region, continued monitoring will be essential for understand the impact on the Ñeembucú Wetlands Complex, a region in which much of the population’s livelihoods depend on ecosystem health. With current ongoing developments in the area and projected continued climatic and anthropogenic pressures, monitoring will be essential for understanding the impact of climate change and development on wetland health. Wetland monitoring is a key tool for addressing wetland change and gaining the knowledge required for well-informed decision making around future development and conservation of valuable ecosystem goods and services.

## Acknowledgements

I would like to thank Fundación Para La Tierra for their support throughout this study and, in particular, to Jorge Ayala, Jack Haines, Sol Gomez, Cristian Torres and Ellen Brogan for logistical and fieldwork assistance. I am grateful to the landowners that gave us permission to survey properties across Ñeembucú.

## Data Availability Statement

The data that support this study will be shared upon reasonable request to the corresponding author.

